# How synchrony and metastable network dynamics are affected in fast and slow timescales with aging: Implication for Cognition

**DOI:** 10.1101/2021.11.29.470424

**Authors:** Priyanka Chakraborty, Shubham Kumar, Amit Naskar, Arpan Banerjee, Dipanjan Roy

## Abstract

Both healthy and pathological aging exhibits gradual deterioration of structure but interestingly in healthy aging adults often maintains a high level of cognitive performance in a variety of cognitively demanding task till late age. What are the relevant network measures that could possibly track these dynamic changes which may be critically relevant for maintenance of cognitive functions through lifespan and how does these measures affected by the specific alterations in underlying anatomical connectivity till day remains an open question. In this work, we propose that whole-brain computational models are required to test the hypothesis that aging affects the brain network dynamics through two highly relevant network measures synchrony and metastability. Since aging entails complex processes involving multiple timescales we test the additional hypothesis that whether these two network measures remain invariant or exhibit different behavior in the fast and slow timescales respectively. The altered global synchrony and metastability with aging can be related to shifts in the dynamic working point of the system based on biophysical parameters e.g., time delay, and inter-areal coupling constrained by the underlying structural connectivity matrix.Using diffusion tensor imaging (DTI) data, we estimate structural connectivity (SC) of individual group of participants and obtain network level synchrony, metastability indexing network dynamics from resting state functional MRI data for both young and elderly participants in the age range of 18-89 years. Subsequently, we simulate a whole-brain Kuramoto model of coupled oscillators with appropriate conduction delay and interareal coupling strength to test the hypothesis of shifting of dynamic working point with age-associated alteration in network dynamics in both neural and ultraslow BOLD signal time scales. Specifically, we investigate the age-associated difference in metastable brain dynamics across large-scale neurocognitive brain networks e.g., salience network (SN), default mode network (DMN), and central executive network (CEN) to test spatio-temporal changes in default to executive coupling hypothesis with age. Interestingly, we find that the metastability of the SN increases substantially with age, whereas the metastability of the CEN and DMN networks do not substantially vary with age suggesting a clear role of conduction delay and global coupling in mediating altered dynamics in these networks. Moreover, our finding suggests that the metastability changes from slow to fast timescales confirming previous findings that variability of brain signals relates differently in slower and faster time scales with aging. However, synchrony remains invariant network measure across timescales and agnostic to the filtering of fast signals. Finally, we demonstrate both numerically and analytically longrange anatomical connections as oppose to shot-range or mid-range connection alterations is responsible for the overall neural difference in large-scale brain network dynamics captured by the network measure metastability. In summary, we propose a theoretical framework providing a systematic account of tracking age-associated variability and synchrony at multiple time scales across lifespan which may pave the way for developing dynamical theories of cognitive aging.

## 1 Introduction

Aging is associated with a gradual deterioration in supporting brain structure [1, 2]. In turn, the alteration in underlying structural connectivity impacts cognitive performances in different domains of cognition including working and episodic memory, processing speed, attention, and a large repertoire of concomitant neurological disorders [3]. The age-related structural decline has been connected with changes in the white-matter fibers, loss of dendritic spines, an increase in number of axons with segmental demyelination, a significant loss of synapses, [4, 5, 6]. Also, the long-range connections specifically undergo age-associated alterations, they themselves act as bridge between distal brain areas, facilitating rapid and efficient interareal communication[7]. One fundamental question that remains poorly understood how does these long rage connections contributes to synchrony and transiently stable brain dynamics at slow and fast time-scales with aging. Here we systematically address this open question to sculpt out a relationship between time scales, anatomical connectivity based on fiber distance and network synchrony and metastability. Leveraging on the empirical observations based on functional magnetic resonance imaging (fMRI) and diffusion tensor imaging (DTI) data allows us to capture the changes in the statistical dependencies of brain signals and their relationship with the change in fiber strength and distance using whole brain computational models in-silico [8].In particular, functional MRI (fMRI) analyses have allowed us to use resting-state scans to better understand the relation between spontaneous fluctuations of the blood-oxygenation level dependent (BOLD) signal (0.1-1.0 Hz) and different resting-state networks (RSNs). Moreover, a discovery of crucial link between age-associated alterations in long-range structural connectivity and network synchronization could provide a mechanistic understanding of neural basis of several existing theories of cognitive aging. Many of the cognitive theories of aging entails interactions and functional coupling between large scale resting-state neurocognitive networks e.g., default (DMN), salience (SN), and central executive network (CEN)[9]. It is found that these networks have shown the most evident and major difference across the young and old age groups [9, 10].

Recent theories further suggest that the dynamical concepts of metastability are suitable for understanding the existing cognitive aging theories[11]. Metastability, a fundamental concept, used to grasp the behavior of complex systems. It is thought to manifest optimal information processing capabilities and switching behavior of the system without becoming locked into fixed interactions [12, 13, 14]. Metastability was also used to confer structure-function integrity in a study by Hellyer et al [15]. They linked reduced metastability of traumatic brain injury (TBI) patients compared to healthy controls due to specific damage to the underlying connectome and correlates well with poorer cognitive flexibility. In the context of aging, metastability has been used to explore the changes in the physiological substrate that take place with aging [11, 16]. Studies using different imaging modalities M/EEG have further showed increased global metastability with age across all known frequency bands of interest [17].

Prior studies suggest BOLD signal variability plays an important role in pinpointing age-associated changes and possibly index cognitive performance of healthy adults [18]. Given the fact that the variability eases flexibility. Garrett et al., further demonstrated both regional increases and decreases of brain signal variability associated with healthy aging has specific functional and behavioral implications [18, 19]. As a consequence, brain signal variability has been widely used to track age-associated shifts and onset of pathological aging process [20, 21]. McIntosh et al., [20] reported using EEG and MEG, with maturation, brain signal variability enhances in the context of local communication between neural populations and decreases for distal communication between neural populations. A widely used measures of metastability (e.g., Kuramoto order parameter) formally capture the variability in the phase synchronization patterns at network level from the BOLD time series data (at slow time scale) or from the EEG and MEG time series data (fast time scale) [22, 23] obtained from specific brain region of interest or at the whole brain level. Hence, we applied the above measure to track age associated shift in metastable brain dynamics and Synchronization at both the time scales of interest to understand the interplay of synchronization among large-scale neurocognitive brain networks. Furthermore, the variability could be critical for the maintenance of cognitive flexibility and performance during healthy aging.

In this work, to address the specific knowledge gap between age associated alterations in longrange structural connectivity and metastable, synchronized interareal brain network dynamics we leverage on whole brain computational modeling. Previous studies have shown how the shift in the working point or demyelination of axonal fiber affects network-level measures [**?**]. Furthermore, since aging entails complex processes involving multiple timescales, therefore, we propose that whole-brain computational models are necessary to test the hypothesis that aging affects the brain network dynamics through network measures of synchrony and metastability at slower and faster time scales. In this study, we use the altered global synchrony and metastability with aging to relate the shifts in the dynamic working point of the system based on two fundamental biophysical parameters e.g., interareal time delay, and interareal coupling strength constrained by the underlying structural connectivity derived from diffusion tensor imaging (DTI) data. We test the auxiliary hypothesis that whether different network measures remain largely invariant or do they exhibit distinct dynamical features in the fast and slow timescales respectively. We simulate a whole-brain Kuramoto model[24] of coupled oscillators introducing appropriate conduction delay and interareal coupling strength to test the hypothesis of shifting of dynamic working points with age-associated alteration in network dynamics in both neural and ultraslow BOLD signal time scales. To achieve realistic brain alteration with age in a model, extremely fine-tuning of parameters is required. So, we tuned the model to resemble empirical restingstate cortical dynamics by evaluating a two-dimensional parameter space, scaling the coupling strength and delay of oscillator interactions. These two biophysical parameters can to capture age-related shifts in parameter space in some studies [25]. This shift might be explained by age-related changes in structural connectivity [26] or loss of a number of long-as well as short-range connections or changes in inter-areal delay due to demyelination [27].

Here, using empirical and computational approaches, we investigate how metastability, defined as the standard deviation of the Kuramoto order parameter [14, 28], arises from underlying structural connectome and is able to track age-related shifts in the dynamic working point. We test 1) how metastability alters globally and locally in resting-state networks (SN, DMN, CEN) with age, 2) whether metastability properties are preserved in slow and fast time scales, and 3) what is the relation between age associated alterations in long-range structural connectivity and metastable, synchronized interareal brain network dynamics.

## 2 Methods

### Subjects

49 healthy subjects (30 females, 19 males; age 18 to 80 years, mean ± SD 41.55 ± 18.44) participated in this study after providing written informed consent. We divided all subjects into two groups comprising 25 young participants ranged in age from 18 to 33 years (mean age = 25.7 ±4 years, 13 female) and 24 elderly participants ranged in age from 55 to 80 years (mean age = 67.99 ± 9 years, 18 female). All experiments were performed in compliance with the relevant laws and institutional guidelines and approved by the ethics committee of the Charité University Berlin.

### Data Acquisition

T1 structural magnetic resonance images (MRI) and diffusion-weighted images (DWI) were acquired at Berlin Center for Advanced Imaging, Charité University Medicine, Berlin, Germany. MRI was performed on a 3T Siemens Trim Trio scanner and a 12 channel Siemens head coil (voxel size). Structural (T1-weighted high-resolution three-dimensional MPRAGE sequence; TR = 1900 ms, TE = 2.52 ms, TI = 900 ms, flip angle = 9./, field of view (FOV) = 256 mm × 256 mm × 192 mm, 256 × 256 × 192 Matrix, 1.0 mm isotropic voxel resolution), diffusion-weighted (T2-weighted sequence; TR = 7500 ms, TE = 86 ms, FOV = 192 mm × 192 mm, 96 × 96 Matrix, 61 slices,2.3 mm isotropic voxel resolution, 64 diffusion directions), and fMRI data (2-dimensional T2-weighted gradient echo planar imaging blood oxygen level-dependent contrast sequence; TR = 1940 ms, TE = 30 ms, flip angle = 78deg, FOV = 192 mm × 192 mm, 3 mm × 3 mm voxel resolution, 3 mm slice thickness, 64 × 64 matrix, 33 slices, 0.51 ms echo spacing, 668 TRs, 7 initial images were acquired and discarded to allow magnetization to reach equilibrium; eyes-closed resting-state) were acquired on a 12-channel Siemens 3 Tesla Trio MRI scanner at the Berlin Center for Advanced Neuroimaging, Berlin, Germany.

### Empirical structural connectivity and tract length data

The empirical structural connectivity (SC) for each subject was generated by using the pipeline described by Schiner et al. [29]. In this pipeline, high-resolution T1 anatomical images were used to create segmentation and parcellation of cortical and sub-cortical gray matter, white matter segments. The main pre-processing steps for T1 anatomical images involved skull stripping, removal of non-brain tissue, brain mask generation, cortical reconstruction, motion correction, intensity normalization, WM, and subcortical segmentation, cortical tessellation generating GM-WM and GM-pia interface surface-triangulations and probabilistic atlas-based cortical and subcortical parcellation. Cortical grey matter parcellation of 34 region of interest(ROI) in each hemisphere was undertaken following Desikan-Killiany parcellation [30]. The connection strength (a value ranging from 0 to 1) between each pair of ROIs was estimated by probabilistic tractography algorithm. SC matrices were generated from each subject’s MRI data and then summed element-wise to obtain an averaged SC matrix.

The pre-processing steps for the diffusion MRI data were eddy current and motion correction with re-orientation of b-vectors (b-zero image was linearly registered to the subject’s anatomical T1-weighted image). Tractography was constrained by seed, target, and stop masks. The fiber length was represented in millimeters. The 68 ROIs or nodes had no self-connection loops meaning that the diagonal values of the SC matrix are all zero.

### Empirical functional connectivity

The same participants were subjected to a functional MRI scan during which their eyes-closed awake resting state data were acquired. The resting-state BOLD activity were recorded for a duration of 22 minutes (TR=2 sec). The BOLD activity was then downsampled to fit the 68 ROIs defined in this parcellation scheme. Aggregated BOLD time series of each region was z-transformed and pairwise Pearson correlation coefficient was computed to obtain the resting state functional connectivity (FC) matrix of each subject. An average 68×68 FC matrix was calculated from each FC matrix. The modeling pipeline was given in Fig. 1 which describes the different stages of the model simulation, as well as, representative results obtained at all different stages of our analysis.

**Figure 1:**
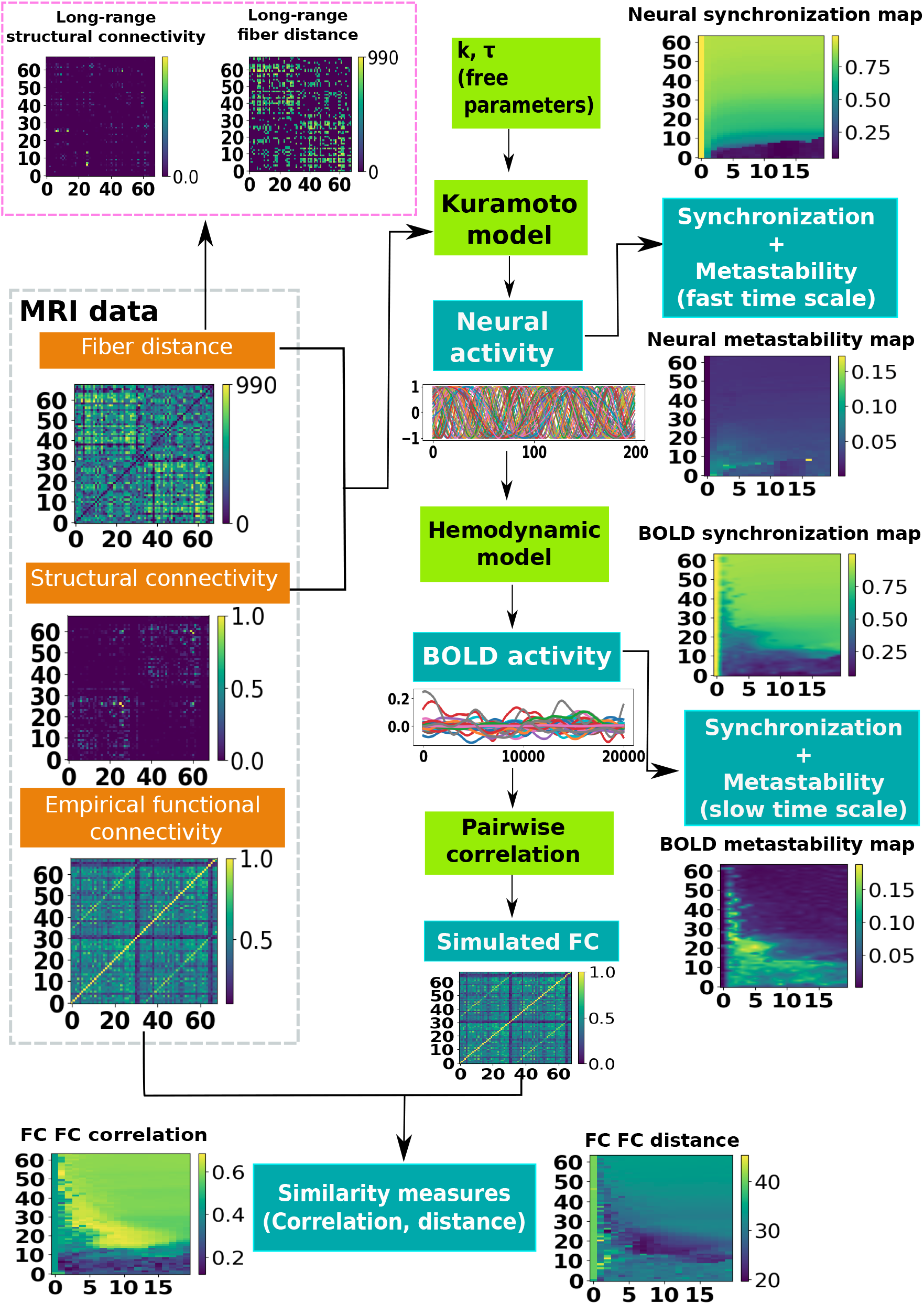
Data driven computational modelling pipeline to simulate whole brain dynamics using the time-delayed Kuramoto model to generate network level synchrony and metastability at slow and fast time scales respectively. Orange text box for experimental data from participants, blue for simulated data, green for generative models to generate network activity at neuronal and BOLD time scales.

### Kuramoto model

In accomplishing our goal, we choose widely accepted Kuramoto model of phase transition and synchronization. As this model with a few anatomical parameters offers tractability of network measures of whole brain synchrony and metastability and also has the potential to capture the key qualitative features of complex brain dynamics enumerated by detailed neural mass models. We considered each *N* (*N* = 68) node as an oscillator and modeled their dynamical interactions using a modified version of the Kuramoto model of coupled oscillators [24, 28, 31]. This version of Kuramoto model takes into account two biophysical parameters, conduction delays and the coupling strength between two nodes. There are two underlying assumptions that local neural activity is periodic and its state can be described by a single variable, the phase. Secondly, there is a weak coupling between local neural populations so that amplitude effects can be neglected [32]. Kuramoto is a less computationally intensive model which can simulate microscopic neural dynamics related to underlying structural connectivity [14, 28, 33, 34].

To address this, the Kuramoto model (with time delay) coupled among *N* brain areas via realistic anatomical connectivity is defined as,

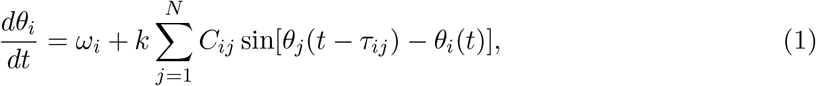

where *i, j* = 1, 2, … *N*. *C*_*ij*_, the asymmetric connectivity (connection weights) between the node *i* and node *j*, normalized to 1. *N* is the total no. of nodes, *θ*_*i*_ is the phase of the node *i, ω*_*i*_ is the intrinsic frequency of oscillation of the node *i, k* is the global coupling coefficient which scales all connections’ strength, and *τ*_*ij*_ is the time delay between the pair of nodes *i* and *j* defined as the ratio of the fiber distance (*L*_*ij*_) and mean conduction speed(*v*), *L*_*ij*_*/v* where *v* is the mean conduction speed of the neural fibers. We can also define a mean delay as *< τ >*=*< L > /v* where *< L >* is the mean fiber length across the brain. We do not consider the effect of noise in this study.

For the sake of simplification, the nodes in the model behave like homogeneous neural masses within which all neurons oscillate together. The intrinsic frequency of oscillation of such neural assemblies has been previously shown to lie in the gamma band [35], and thus in our model we fix the value of *ω* = 2*π* × 60 Hz. The nodes are free to oscillate, with the phase of the oscillation, *θ*, defining the state of the node. This network is thus a phase oscillator. The free parameters in this model are *k* and *< τ >*(ms), for which we performed parameter space sweeps in order to characterize the network dynamics for the entire set of relevant parameter ranges, and the plots so computed are called the parameter maps.

Random initial conditions were selected for simulating phase evolution with respect to time. Furthermore, as this model incorporates time delay we had to specify the phases for a sufficiently long time interval, say 80sec in order to capture the steady state dynamics and first 10sec were discarded for all subsequent analysis. The system of *N* dynamical equations was numerically integrated by using a variant of the Euler method adapted to noise with a time step Δ*t* = 0.1 ms. All calculations were performed in MATLAB 2016b (MathWorks) and BrainNet viewer and Python 3.8 were used for plotting the figures.

### Synchronization and metastability

The dynamics of the Kuramoto network of oscillators can be characterized by the order parameter *R*(*t*), which is defined below,

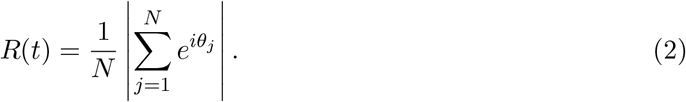

*R*(*t*), the coherence of oscillators at time *t*, ranges from 0 for a fully desynchronized or incoherent state to 1 for a fully synchronized state.

To derive empirical metastability, we first calculated the analytic signal (*y*_*j*_) for BOLD signals from each brain regions of interest as follows:

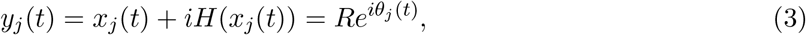

where *H*(*x*_*j*_(*t*)) represents Hilbert transform of the original signal *x*_*j*_(*t*) at *j*-th ROI. Analytic signal has advantage over original signal since it discards negative frequency components without loss of information and makes instantaneous phase (*θ*(*t*) of the signal accessible; hence allowing to explore relationships at higher temporal resolution. Kuramoto order parameter defines mean phase synchronization or instantaneous coherence in the network as

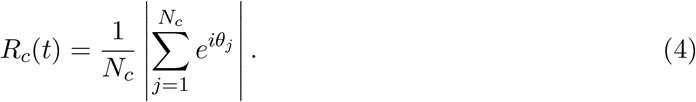

Here, *N*_*c*_ represents the number of regions (node1s) in th1e network *c*. For whole-brain analysis, *N*_*c*_ = 68, and for different large scale resting-state networks *N*_*c*_ depends on the number of regions considered. Then metastability is defined as standard deviation of *R*_*c*_(*t*) over time.

Fo synchronization at frequency Ω, equation 1 reads [36],

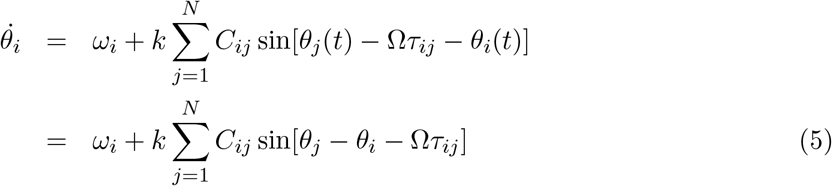

The phase difference between each pair of Kuramoto oscillators is given as,

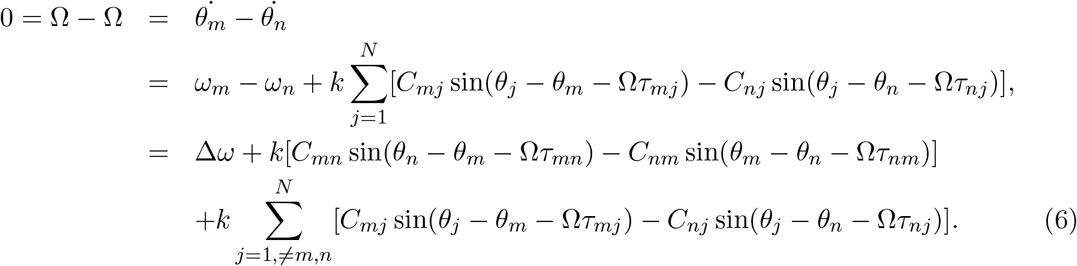

For symmetric coupling *C*_*mn*_ = *C*_*nm*_ = *c*(say), *τ*_*mn*_ = *τ*_*nm*_ = *τ* (say). In our experiment Δ*ω* = 0. Let, Δ*θ* = *θ*_*m*_ − *θ*_*n*_, then equation 6 reduced to

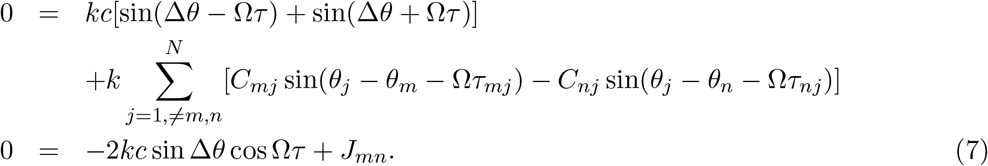

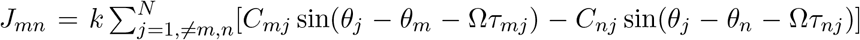 contains all the other links towards the nodes *m* and *n* apart their direct link. Therefore,

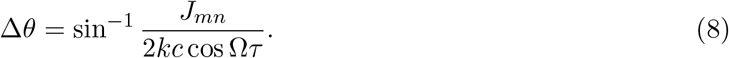

For synchronized state, Δ*θ* = 0, above equation reduced to

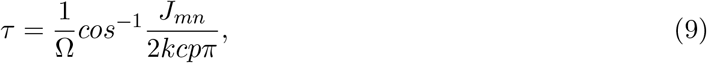

where *p* ∈ ℕ.

### Simulated functional connectivity

The simulated neural activity from the Kuramoto model was converted to the BOLD activity to have a direct correlation with the empirically recorded BOLD activity from the subjects. The simulated neural fluctuations were given by the firing rate *r*_*n*_(*t*) for node *n* fluctuates around a fixed value and these fluctuations were obtained by a periodic function of local node phase [28]. We chose a simple sine function for these fluctuations as *r*_*n*_(*t*) = *r*_0_ sin(*θ*_*n*_(*t*)). For simplicity, we considered the amplitude of the model as *r*_0_ = 1. Then we used the Balloon-Windkessel hemodynamic model to convert the neural signal to BOLD activity. The method was adopted from previously published works [28]. Snapshots of neural activity for a particular parameter set (*k, τ*) at both these levels were shown in Fig. 1 which were seen to be widely variant from each other.

The simulated BOLD activity was then pairwise correlated to obtain the simulated FC. We calculated the Pearson correlation and the distance between the two matrices only for the structurally connected pairs to compare the simulated and the empirical FCs. In Fig. 1, we plotted the measures of similarity between the empirical and simulated FC, identify the regions of the parameter space where the model predict the experimentally observed functional connectivity to a remarkable degree. In the central yellow region of the parameter map of FC-correlation, with values around 0.6 the model provides the best fitting results. Hence, in this regime in the parameter space, this model could be used as convenient source for generating realistic neural activity. The parameter map of Euclidean distance also identified the region around the best fitting values, shown as the dark-blue region. As expected the regions thus identified by the two independent similarity measures show significant overlap and unless otherwise specified is used for the remaining analysis.

### Selection of brain regions from large scale brain networks

We considered three resting-state large scale brain networks, namely the SN, CEN, DMN (node details were provided in Table 1). Several studies reported the alteration of interconnections within and between the DMN, CEN, SN in many psychiatric and neurological disorders, for instance, Alzheimer’s disease, psychosis, attention deficit/hyperactivity disorder, autism spectrum disorder, and depression [37]. DMN comprise of inferior parietal lobule (IPL), posterior cingulate cortex (PCC), and medial orbitofrontal cortex (MOF) [38]. These regions are considered as ‘default mode’ of brain function, as they exhibit decreases in activity during a variety of goal-directed behaviors. Bilateral rostral and caudal middle frontal gyrus (MFG) and superior parietal lobule (SPL) were selected as nodes of CEN [39]. It plays an important role in decision making and executive functions. It has been consistently reported that DMN and CEN maintain antagonistic relationship in resting state or during task performing in healthy individuals [40]. SN, which comprises of anterior insula and caudal, rostral ACC was previously used in our study [9], is important for the detection of salient events and switching between other large-scale brain networks in resting as well as task conditions [41, 42]. SN enables rapid recognition between goal-directed or goal-oriented salient stimulus from external environments and plays important role in the dynamic switching of antagonistic activity between the DMN and CEN. But how this dynamic switching alters in normal aging is unknown.

**Table 1:** Coordinates of selected nodes of three resting-state networks according to Desikan–Killiany (DK) parcellation atlas.

| Networks | Brain regions | MNI coordinates(x,y,z) |  |
| --- | --- | --- | --- |
|  |  | Left(l) | right(r) |
| Salience network | Insula | (−41, 13, −6) | (43, 12, −6) |
|  | CACC | (−2, 21, 27) | (3, 21, 27) |
|  | RACC | (−2, 39, 6) | (4, 38, 4) |
| Central executive network | RMFG | (−34, 53, 17) | (43, 45, 21) |
|  | CMFG | (−45, 18, 46) | (43, 14, 43) |
|  | SPL | (−25, −62, 63) | (17, −65, 59) |
| Default mode network | MOF | (−4, 44, −14) | (7, 45, −13) |
|  | IPL | (−47, −70, 31) | (48, −67, 29) |
|  | PCC | (−1, −18, 38) | (1, −16, 37) |

### Determination of long- and short-range connections

Structural connectivity is described by fiber tracts joining different brain regions at the macro scale [43].dMRI-based tractography is used to characterized SC properties e.g., fiber length and strength. There have been numerous definitions of long- and short -range SC in the extant literature [44, 45] depending on their tract lengths. In this work, we used fiber tract length to define long- and short-range connections in the following manner. In Fig. 2(A), we plotted the average fiber tract lengths between each pair of regions for the young group. Longer connections between two regions were depicted by thicker edges whereas thinner edges indicate short-range connections. Distribution of average fiber tract length for young group has been plotted in Fig. 2(B). First quartiles, (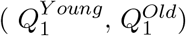) and third quartiles, (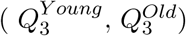) were calculated from average fiber tract for both young and old group. Subsequently, we considered the minimum value of first quartiles 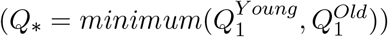 as lower threshold and maximum value of third quartiles 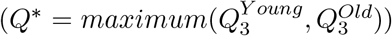 as upper threshold. Now, the tract lengths higher than the upper threshold considered as long-range connections and the tract lengths lesser than the lower threshold considered as short-range connections. The blue vertical lines in Fig. 2(B) indicate the lower threshold (*Q*_*\**_ = 222mm) and upper threshold (*Q*^*\**^ = 581mm) respectively. The threshold matrices were shown in Fig. 2(C,D). Fig. 2(C) showed only the long-range connections and Fig. 2(D) showed the short-range connections for young cohort.

**Figure 2:**
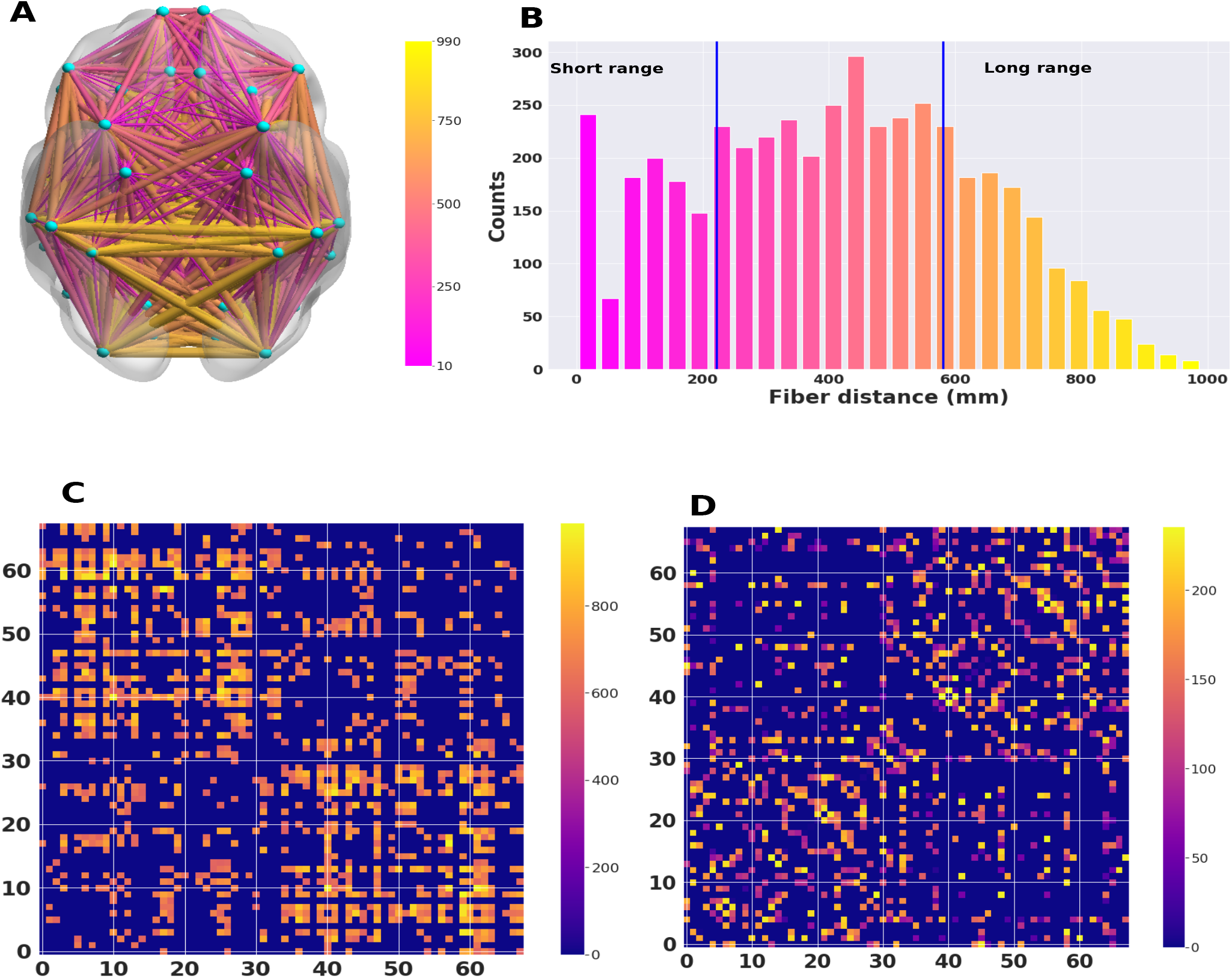
Determination of long- and short-range white matter tracts. (A) Schematic illustrating connection length profiles between each brain regions. (B) Based on the distribution of the tract lengths in young cohort, structural connections were categorized into short-range (*<* 222 mm) and long-range connections (*>* 581 mm). Average (C) long- and (D) short-range connections in all 68 brain regions for the young cohort.

We have deleted top 5% long-range structural connections from structural connectivity matrix of young subject and simulated the model keeping other parameters fixed. Then the difference was calculated by subtracting the global metastability of young subjects from modified global metastability with deleted connections. We repeated the calculation of difference of global metastability after removing 10%, 15%, and 20% long-range connections.

For similarity between structural connectivity matrices, we used Kolmogorov-Smirnov (KS) similarity as 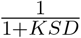, where *KSD* > 0 is the Kolmogorov-Smirnov distance between two matrices.

### Statistical analyses

We have applied Mann Whitney U test to compare group differences between young and old subjects. In addition, the two sample Kolmogorov-Smirnov (KS) test was conducted to investigate whether long-range and short-range connections of older subjects were more disrupted than long-range and short-range connections of younger subjects. KS tests a null hypothesis by comparing whether two groups are from the same populations with identical distributions or not. It does not compare any particular group statistical average quantities e.g., mean or median of a distribution. For group comparisons, *p*-values of *<* 0.05 were considered statistically significant after FDR corrections.

## 3 Results

### Empirical measures of metastability for large-scale brain network dynamics

As discusses in the previous section, network level synchronization among the large-scale brain networks were conveniently captured by the Kuramoto order parameter, *R*(*t*). The order parameter *R*(*t*) exhibiting synchronization dynamics as a function of time is plotted for the young and older group in Fig. 3(A) for the whole brain network (defined as global) and for the three large-scale brain networks of interest as shown in Fig. 3(B). Interestingly, we found that the younger group showed higher values for *R*(*t*) while maintaining lesser fluctuations than the order parameter dynamics for the older group. Next, we assessed the metastability of largescale neural dynamics, which was measured using 68 regional phase time courses obtained from resting-state fMRI BOLD data in both young and old subjects and plotted in Fig. 3(C). Then, we compared metastability within resting-state networks between young and old by using Mann Whitney U test. Global metastability increased with age but not significantly. Old subjects showed significantly greater metastability in SN (*U* = 206, *p* = 0.04). Metastability in CEN and DMN increased for older population but not significantly. However, findings for empirical metastability did not survive false discovery rate (FDR) corrections.

**Figure 3:**
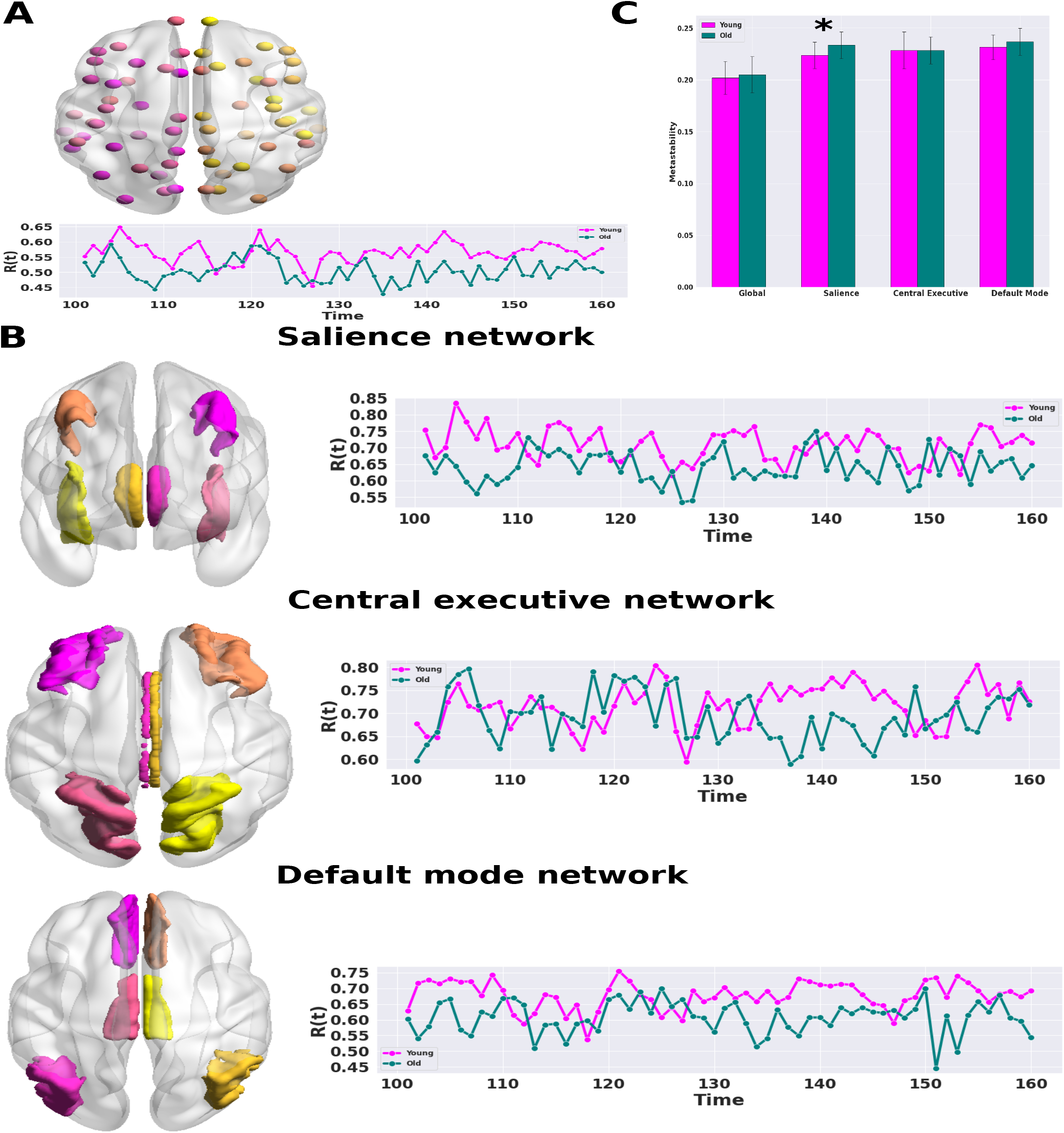
Global and local order parameters and empirical metastability for young and old cohorts. (A) Spatial node map of all the regions and corresponding order parameter, *R*(*t*) for both groups. (B) Spatial ROI maps and order parameter, *R*(*t*) of the three resting-state networks as salience, central executive, and default mode network respectively for both young and old cohort.(C) Bar plot of empirical metastability of young and old subjects. Empirical metastability at rest was increased in older subjects compared with younger subjects; Mean measures of metastability (± SD) estimated using a phase-transformed functional time course extracted from 25 young and 24 old subjects suggest that global measures of metastability are increased with age, significantly for salience network (“*” signifies *p <* .05, not FDR corrected) and not significantly for the global, central executive network and for default mode network. Colors of the nodes and ROIs are not signifying any values.

### Generation of cortical activity from the Kuramoto model

We used a generalized Kuramoto model with time delay for the generation of cortical activity, following the procedure described previously in [28] among others [46, 47]. The input parameters in this modeling scheme were data from MRI diffusion imaging and fiber tractography studies, i.e. the structural connectome and the fiber distance information termed as the structural connectivity (SC) matrix and distance matrix respectively. The global dynamical behaviors of the model, synchronization and metastability are characterized by the mean synchronization level *R* and standard deviation of *R*(*t*) over the simulated time interval, respectively (Fig. 4). Fig. 4 (A) showed the behavior of the Kuramoto model at *t* = 2000ms for three different values of coupling strength keeping other parameters fixed. At low coupling strength (*k* = 5) (Fig. 4 (A) left) each node behaved incoherently, then for *k* = 27 (Fig. 4 (A) middle) nodes were partially synchronized, and for *k* = 35 (Fig. 4 (A) right) nodes were mostly synchronized. In Fig. 4 (B), the order parameter *R*(*t*) (red line) with metastability (yellow confidence interval) were plotted for three different coupling strengths as defined in Fig. 4(A). Fig. 4 (C) showed metastability was very low if the system is either completely synchronized or completely de-synchronized and a large value indicated switching between coherent and incoherent states.

**Figure 4:**
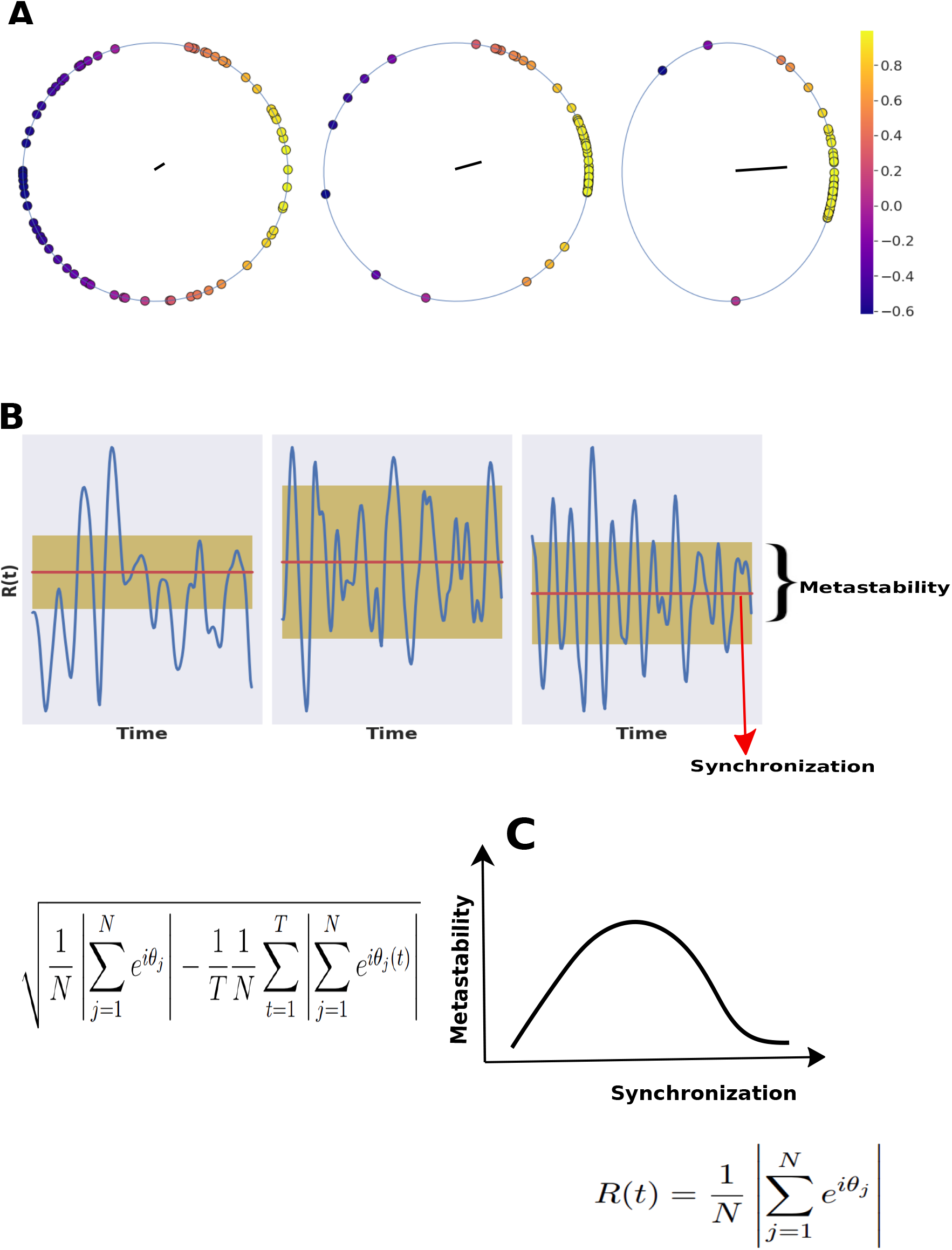
Schematic depicting phase synchronization and metastability. (A) Color code phase circle diagram for desynchronized (left), partial synchronized (middle), and synchronized state(right); (B) Order parameter (R(t)) for three different nature of synchronization, the red line is global synchrony (mean of *R*(*t*)), and the yellow confidence interval is metastability (standard deviation of R(t)); (C) Relationship between synchronization and metastability, metastability is maximum when the system is partially synchronized.

The similarity between the simulated FC and the empirical FC from fMRI data was measured by taking the Pearson correlation (Fig. 5(A, B)) and Euclidean distance (Fig. 5 (C, D)) plotted in the two dimensional parameter space given by (*k, τ*). It was immediately noticeable that the young group in this data exhibited regions of high correlation, with values as high as 0.6 highlighted by the contour lines in Fig. 5 (A). In contrast, the correlation in the old group, as shown in Fig. 5 (B), was relatively lower along with the island of high correlation being substantially reduced. The FC Euclidean distance measure, on the other hand, for the two groups have a larger overlapping area of agreement in the parameter space. The region of simultaneous large correlation and small distance corresponds to a mean delay between 12ms and 24 ms and inter-areal coupling strength value between 15 to 25 for young population and mean delay of 15ms to 22ms along with coupling strength of 12 to 20 for the older population.

**Figure 5:**
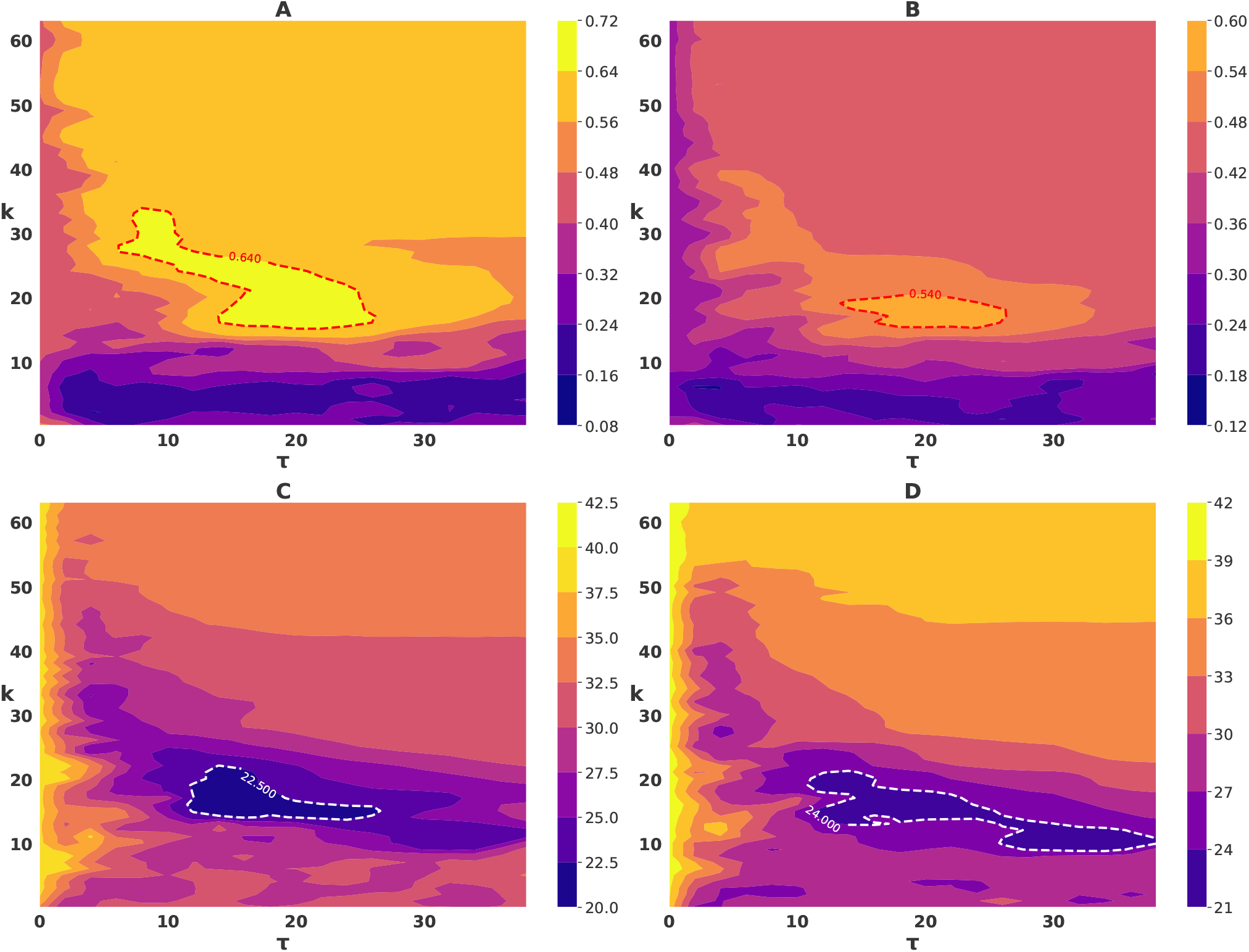
Contour plot of FC-FC correlation and Euclidean distance between empirical and simulated FC for young and old subjects in (*τ*, k) parameter space. The plots show: (A,B) FC-FC correlation for young and old subjects respectively. The red dotted lines correspond to the highest FC-FC correlation values. (C,D) FC-FC Euclidean distance for the same. The white dotted lines correspond to the lowest FC-FC distance values. The region of high correlation was confined to a much smaller area with smaller correlation value in the case of the older population group.

### Synchronization and metastability in slow and fast time scale

Next, we simulated the whole brain dynamics in the best fitted (*τ*, k) parameter space to test whether model predicted network synchrony and metastability associated with age is somewhat comparable against metastability results obtained from empirical observations. We evaluated synchrony and metastability globally in the (*τ*, k) parameter space for BOLD time signals after discarding initial 10 seconds of transients. The parameter maps of BOLD signal synchronization were depicted in Fig. 6 (A,B) and similarly, metastability were plotted in Fig. 6 (C, D) for both age groups. The older group displayed slightly higher synchronization values than the younger cohort. However, the parameter maps of metastability (Fig. 6 (C, D)) complement the results of the synchronization maps, where regions of high metastability coincides with the part of the parameter regions where partial synchronization was manifested. Interestingly, there was not much difference in the whole brain metastability between the two groups suggesting some kind of invariance and preservation of metastable brain dynamics associated with healthy aging process.

**Figure 6:**
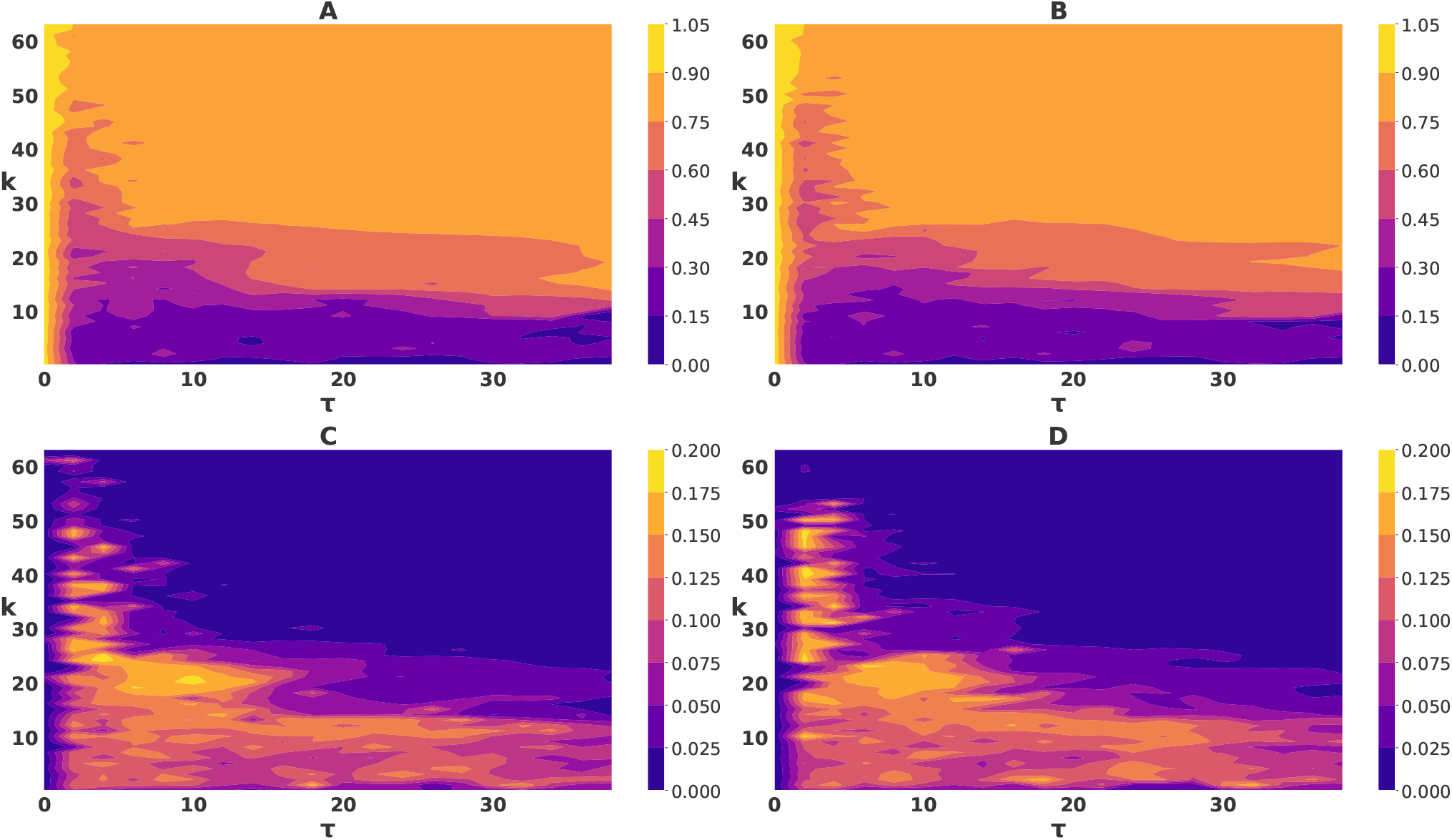
Global dynamics in the mean delay and global coupling parameter space in BOLD time scale. Profiles of the synchronization in (A) young, (B) old cohort, and profile of metastability in (C) young, (D) old cohort.

The dynamics of simulated BOLD signals (for all 68 nodes) based on different combinations of (*τ, k*) were plotted in Fig. 7 to demonstrate rich network dynamics comprise of Synchrony, Partial Synchrony, Incoherence. To systematically explore the emergent network dynamics we have selected three sets of parameters respectively (*τ* = 8, *k* = 5), (*τ* = 20, *k* = 10), and (*τ* = 12, *k* = 35) from the parameter space where synchronization was low, moderate and high respectively and plotted the corresponding dynamics in Fig. 7 (A,B), Fig. 7 (C,D), and Fig. 7 (E,F) respectively. It was clear from the Fig. 7 (A,B) that phases of the signals were mainly incoherent whereas the phases of the signals in Fig. 7 (E,F) were synchronized. But in Fig. 7 (C, D) some phases were synchronized and some of them were desynchronized.

**Figure 7:**
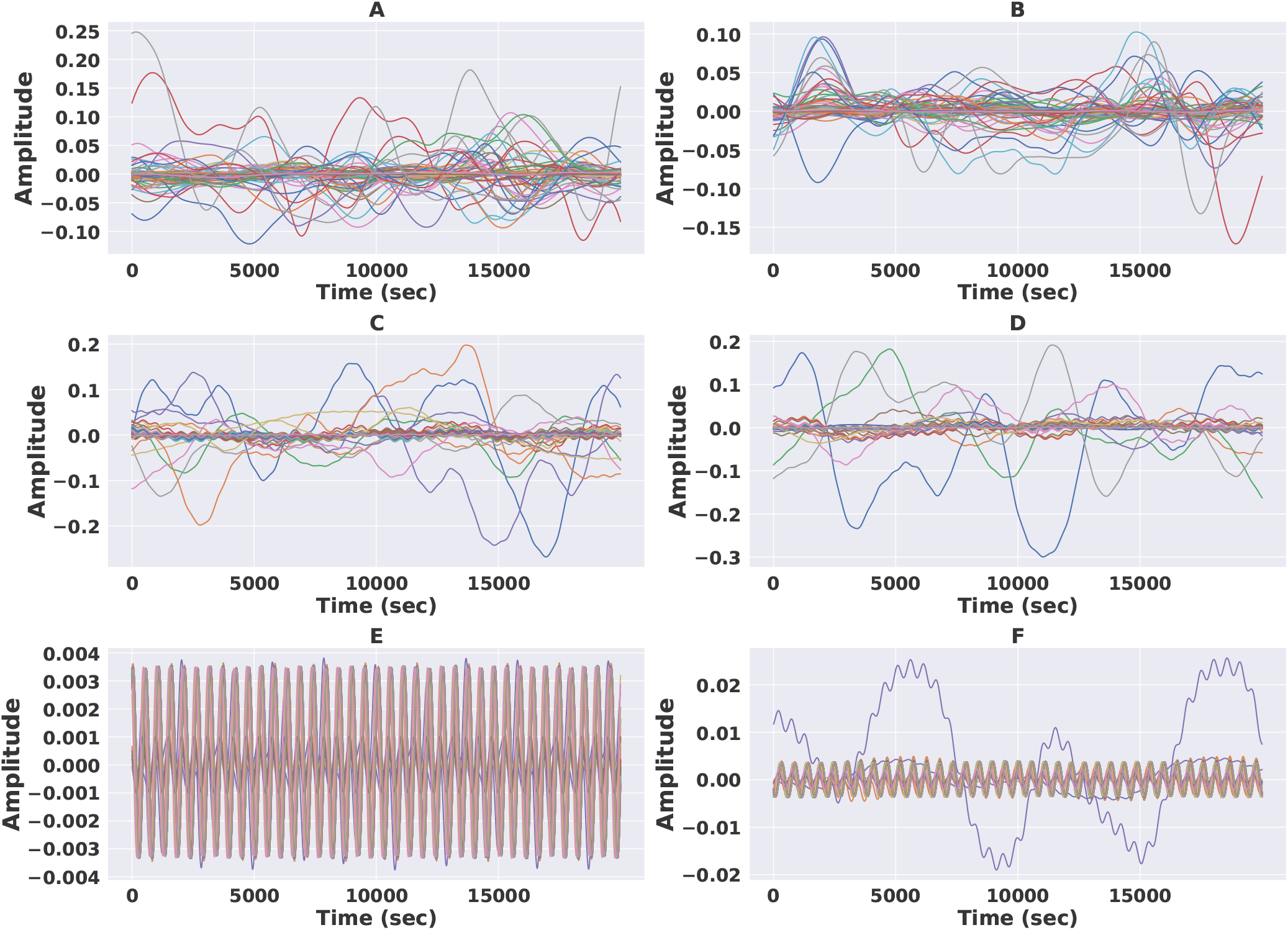
BOLD time signals for young group (left column) and old group (right column) at different points in parameter space. The parameter values are (A,B) *τ* = 8, *k* = 5; (C,D) *τ* = 20, *k* = 10; and (E,F) *τ* = 12, *k* = 35.

Next, we will focus our interest on the fast time scale, also known as the neural time scale, to understand our auxiliary research question whether the network measures Synchrony and metastability patterns remains largely invariant or distinguishable across time scales with healthy aging. As a first approximation, the fast time scale brain network dynamics is simply the time evolution of the phases of the individual oscillators. As we are mainly interested in the synchronization and metastability of the brain, we calculated these values, and performed a parameter sweep across the two variables *τ* and *k*, in order to get the parameter maps of synchronization and metastability. Fig. 8 (A,B) depict the parameter space of network synchronization and Fig. 8 (C,D) showed the parameter space of metastability for both groups at the neuronal time scale. While the network synchronization were nearly identical for both age groups the network metastability was slightly higher in the older group. The parameter maps of synchronization, in Fig. 8 (A,B) for both age groups, showed with narrow contour lines (marked in white) where rich and complex brain dynamics co-exist. In this regime, the network was neither in a completely phase locked state (yellow regions) nor in decoupled incoherent state (blue regions). This region was wider for older group than younger group suggesting aging probably shifts the identified parameter space and supports higher transient synchronization. Interestingly, it can be further observed from Fig. 8(C,D) that the higher values of metastability lied in the same regions of the parameter where partial synchronization occurs. This is quite similar to the observations made on the slower time scale. This further crucially implies metastable brain dynamics is not only preserved across healthy aging individuals but also across two time scales of information processing and cognition.

**Figure 8:**
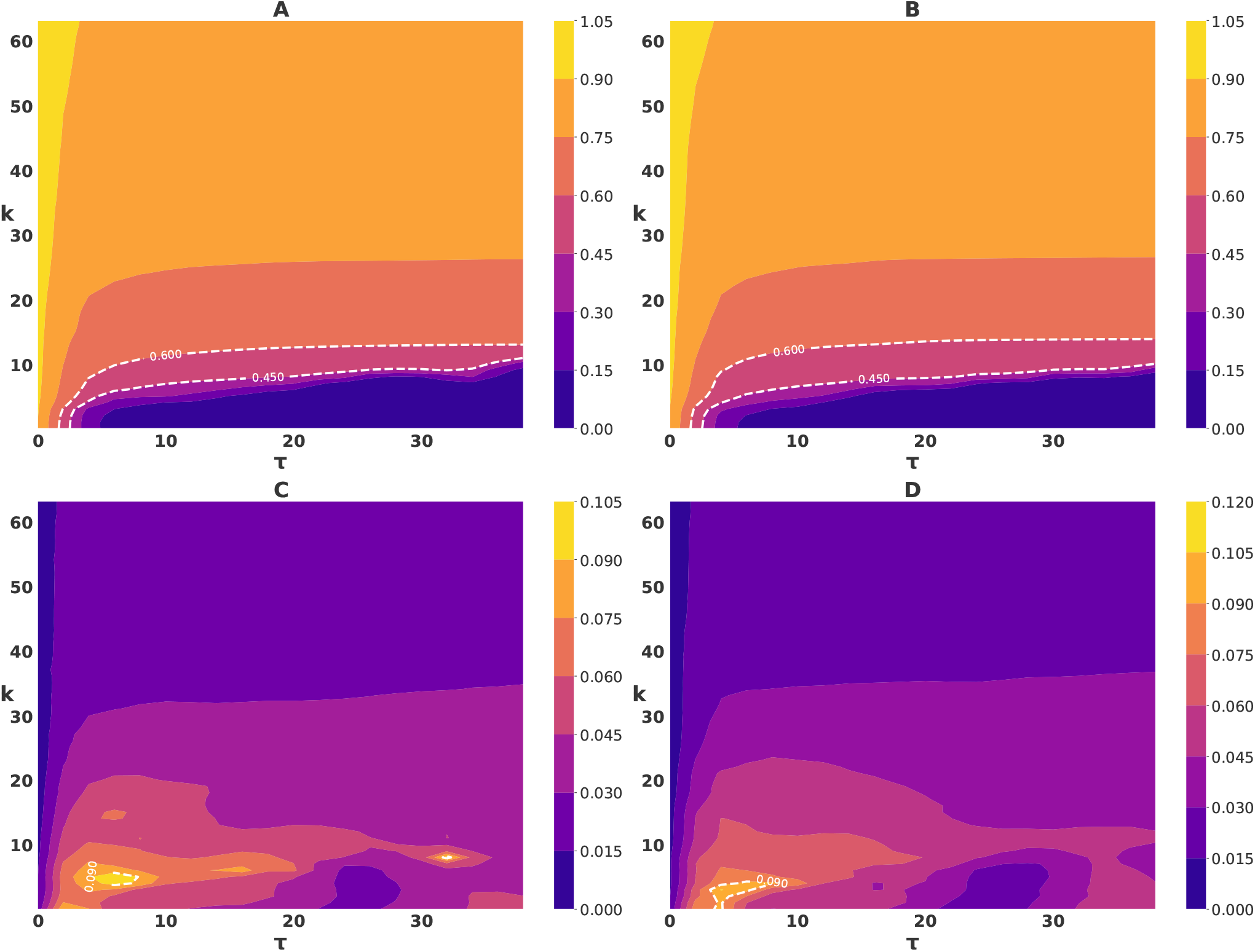
Global dynamics in the mean delay and global coupling parameter space in neural time scale. Profiles of the synchronization in (A) young, (B) old cohort, and profile of metastability in (C) young,(D) old cohort.

The dynamics of the neural oscillations across all the 68 cortical regions in the young population were represented in the plots of Fig. 9 for different combination of parameters representing different dynamical regimes as desynchronized, partially synchronized, and fully synchronized respectively. These neural signals were only a portion of the total time of activity. Parameters *τ* = 8, *k* = 5 lied in desynchronized regime so the neural signals were desynchronized as we can seen from Fig. 9 (A,B). Similarly,the parameters, (12, 20), (0.5, 40) were in partially synchronized and synchronized regime and from Fig. 9(C,D) and Fig. 9(E,F), we can verify the natures of the oscillations.

**Figure 9:**
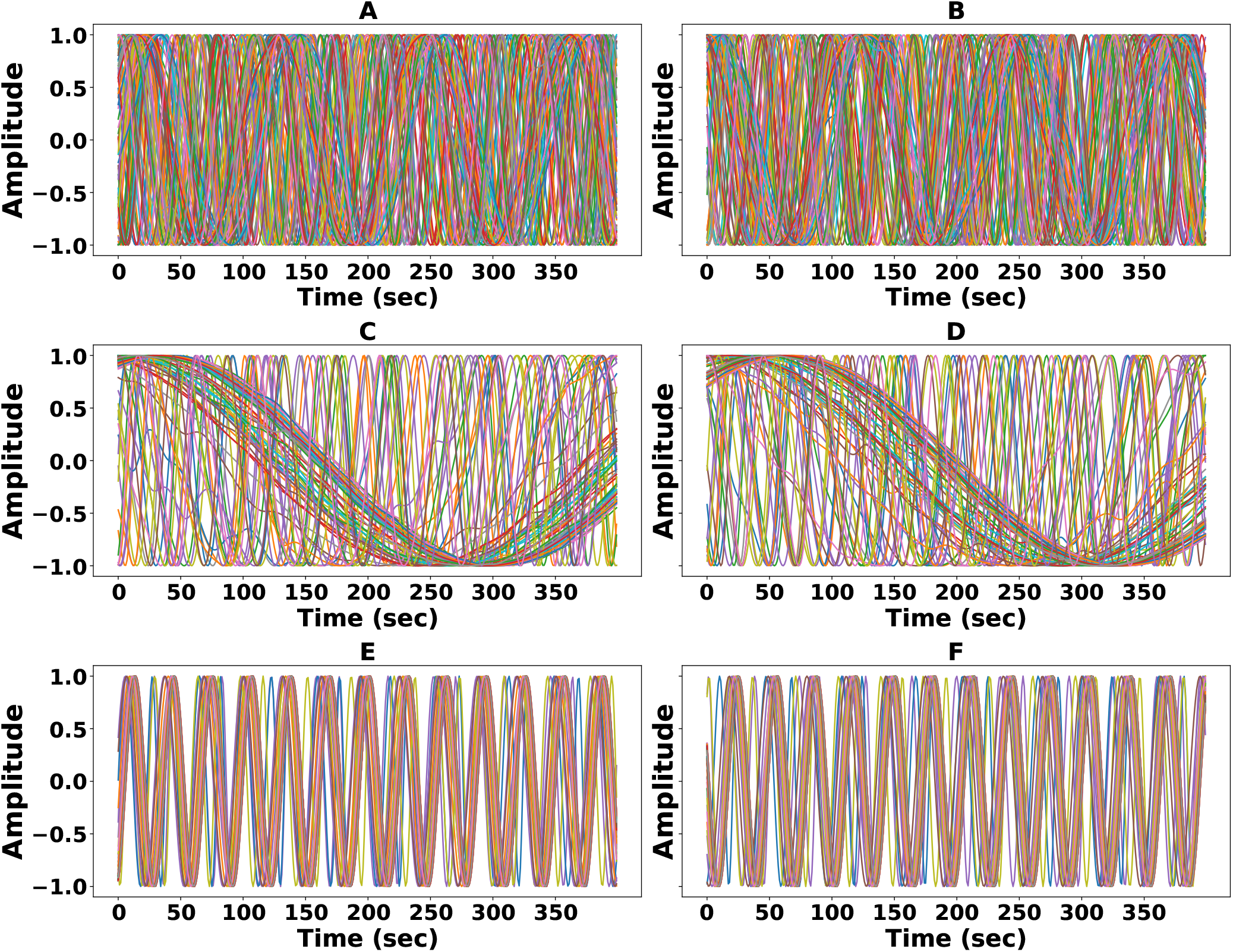
Neural time signals for young group (left column) and old group (right column) at different points in parameter space. The parameter values are (A,B) *τ* = 8, *k* = 5, (C,D) *τ* = 12, *k* = 20, and (E,F) *τ* = .5, *k* = 40. For, (A,B) all the signals are mostly desynchronized; (C,D) the signals of some regions are synchronized whereas some are desynchronized; and (E,F) most of the signals are synchronized.

### Metastability of resting-state functional networks

Next, we analyzed the metastability of resting-state networks (RSNs), crucially, the three large-scale neurocognitive networks defined earlier, the salience (SN), central executive (CEN), and default mode network (DMN). The results presented in Fig.10 showed a comparative study of the variation of metastability across the two age groups, for these three different functional networks SN, CEN and DMN. Fig. 10 (A) depicted the BOLD metastability map in SN, CEN and DMN for young and old group. While the metastability of the CEN and DMN do not show major variations with the age, we found that the metastability of the SN increased slightly with age. This can be seen in the higher metastability values, spread over a larger area of the parameter space in the case of the older population. Moreover, the increased metastability, visualized in the parameter space, occurred in the region where the parameters were relevant to realistic values and within which the model had high prediction accuracy. The results of the empirically obtained metastability at the BOLD timescale was compared to the simulated metastability across the entire parameter space. As a consequence of parameter search, we have identified regions in the parameter space where the trend of the metastability with age either agrees or disagrees with the empirically observed results. This grid search procedure further allowed us to constrain and predict biologically relevant (*τ, k*) values. For those values of parameters where the metastability of all the four cases (global (whole brain), SN, CEN, and DMN) showed an increase with age, agreeing with the empirical observation, we denoted those points on the parameter space by a circle (o). On the other hand, for the critical parameter values those resulted in disagreement with the experimental results, denoted by a cross(x). In this way, the entire parameter space was characterized and the results summarized in Fig. 10 (B) with representative examples being shown in Fig. 10 (C) for BOLD time scale and in Fig. 10 (D) for neural time scale respectively. This helped us identify the areas of the parameter maps where the simulated results and empirical results were in qualitative agreement.

**Figure 10:**
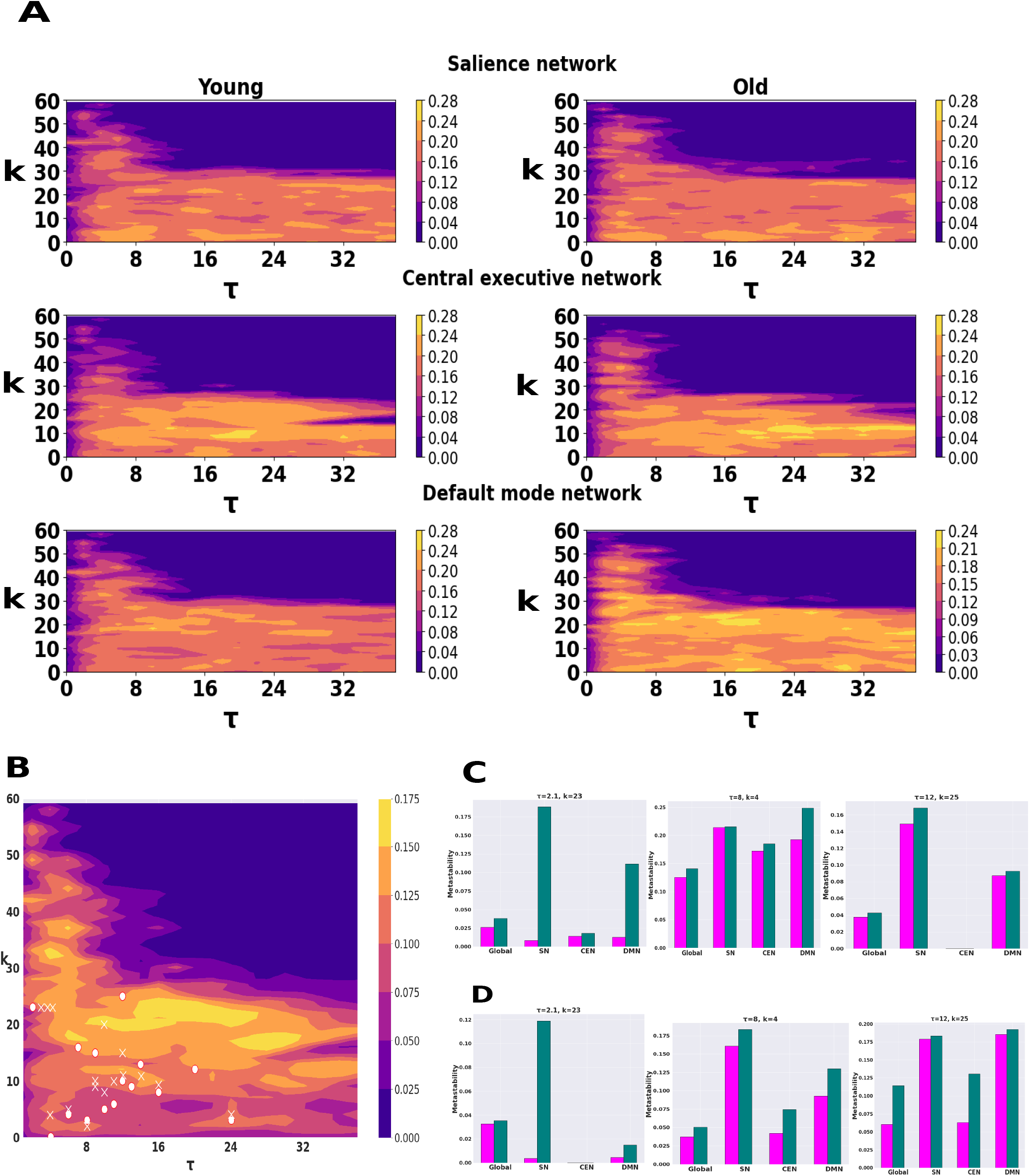
(A) A comparison of the parameter maps of metastability for the SN, CEN, and DMN in the young and old cohorts shows a marked increase in the metastability of the SN for the old cohort, but not so in the case of the CEN and DMN. (B) A summary of the different basins of attraction in the parameter space, showing regions of agreement (o) and disagreement (x) of the empirical and simulated metastability. (C) Representative examples of metastability of BOLD signals for both populations at different points on the parameter space marked by (o). (D) Metastability of neural signals for same points defined in (C).

### Analysis of structure

Next, we hypothesize that the observed differences in the metastability for the young and the old groups across different neurocognitive networks may be arising from the difference in the proportion of short-and-long-range connectivity in the underlying SC matrix. Therefore, we attempted to analyze the differences in the structural connectivity patterns across these two groups. The SC weight versus fiber length distribution may be visualized by plotting the SC edge strength with respect to the fiber lengths. In the human brain, and indeed in other species as well, this relation follows a somewhat linear profile when plotted in the log scale for the edge strength [48]. We plotted the SC weight versus fiber length distribution for the brains of a young and an old subject in Fig. 11 (A). We used the slopes of these distributions for each subjects, obtained by a linear regression, as the marker for characterizing the particular network. Mean slopes for both groups in global and in three networks (SN, CEN, and DMN) were plotted in Fig. 11(B). Non parametric, Mann Whitney U test has been applied on slopes and no significant differences have been found for global (*U* = 257, *p* = 0.14), for salience (*U* = 309, *p* = 0.48), for central executive (*U* = 288, *p* = 0.32). Only default mode network (*U* = 214, *p* = 0.04) showed significant decrease of slopes for older individuals. But this effect did not survive FDR correction. This means that the long-distance fibers of DMN in older subjects had very smaller connection strength than that of younger subjects. We did not see any major differences in the distribution profiles (slopes) of the brains of the young and the old groups.

**Figure 11:**
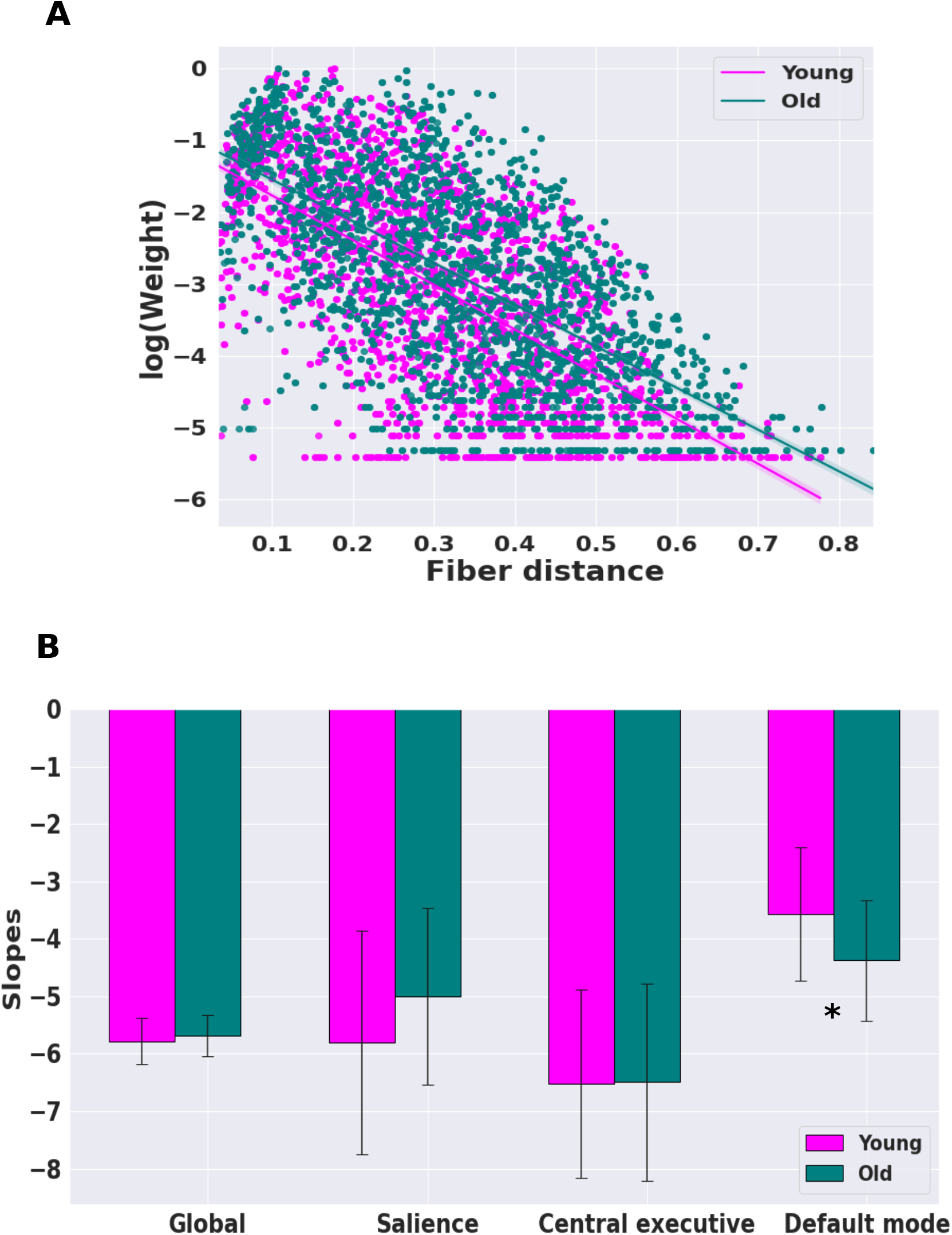
(A) A comparison of *log*(SC) weight vs normalized fiber distance distributions for the whole brain for a young and old subject. Regression lines are fitted to individual scatter plots to find their slopes and intercepts of the distributions. (B) Summary of the mean slopes of the SC weight-fiber distance distributions for the whole brain, salience, central executive, and default mode network for the two groups. From Mann Whitney U test, we found that the mean slope of older subjects was significantly lesser than slope of younger subjects for the default mode network. “ * “ signifies *p <* .05(Not FDR corrected).

Now, in Fig. 12, we explored the changes in long- and short-range connections with age. Violin plots were drawn in Fig. 12 (A) for long-range connections and Fig. 12 (B) for short-range connections of young and old subjects. We applied the KS test to compare whether these distributions of long-range counts as well as short-range counts were significantly different or not. The number of long-range connections was significantly reduced for the old group (*p <* 10^*−*5^, FDR corrected), though the median values of the long-range fiber tract were almost remain same as 683.94 for young and 660.61 for old group. The counts of short-range connections were reduced for the older population but not as significant as the counts long-range connections. The histogram plot of long-range connections for both groups together was shown in Fig. 12(C). The dotted lines represent the kernel density estimation for individual distributions. So many long-range counts were lessened for older population. By Fig. 12(D), vertical distance between the two cumulative frequencies of long-range connections of young and old groups were increasing after tract length of 700 mm. Finally, we plotted in Fig. 12(E), the circular maps of top 10% long-range connections for both groups and found that the stronger connections were mainly damaged by age. So the length of long-range fiber tract was remain unchanged whereas the number of long-range connections were significantly decreased with age.

**Figure 12:**
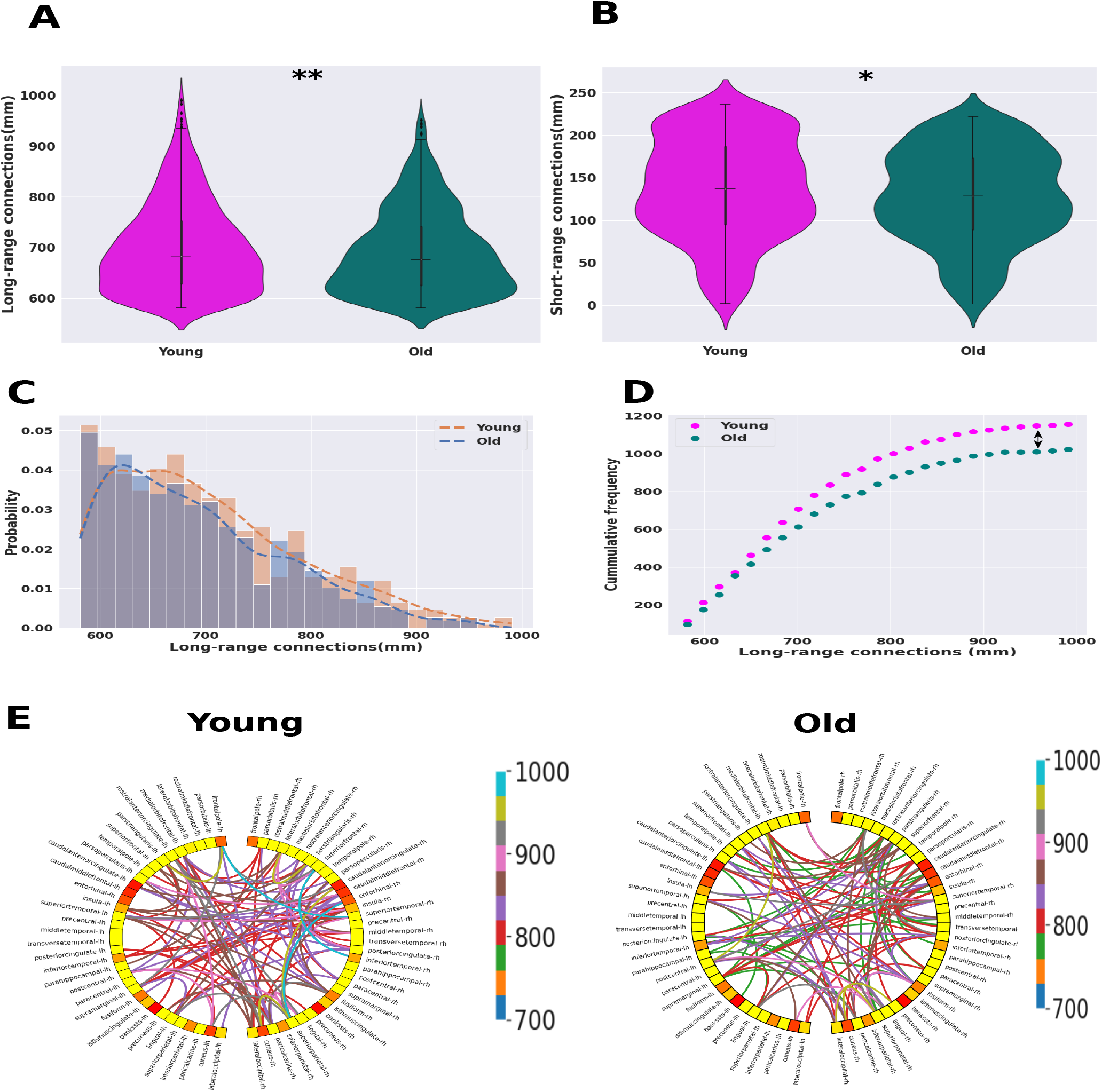
Violin plots of (A) long-range and (B) short-range fiber tract lengths for young and old groups. The white dot represents the median value,the thick gray bar in the center represents the interquartile range. Two sample KS test had been performed to compare two distributions. The counts of long-range connections were significantly reduced for the older population. The counts of short-range connections were reduced for the older population but not as significant as the counts long-range connections. (C) Histogram plot of long-range connections for young and old groups. The dotted lines represent the kernel density estimation for individual distributions. (D) Cumulative frequency counts of long-range connections for younger and older individuals. Vertical arrow line in the plot represents the KS distance. As the length of long-range connections increase, the number of connections decrease for the older group than the younger group. (E) Circular graph plots of top 10% long-range connections for both groups. So, the numbers of stronger long-range connection were damaged with age. *p <* 10^*™*5^ is indicated by “ **” and “ *” signifies *p <* .001, FDR corrected.

Next, we computed the KS similarity between modified structural connectivity matrix of young subject with 5% deleted long-/short-range connections and structural connectivity matrix of old subject. We did this for various percentage of deleted connections and plotted the similarity values in Fig. 13(A). It is clear that when we deleted the long-range connections from young subject’s structural connectivity, the new structural connectivity matrices were highly similar to structural connectivity matrix of old subject. If we compare the similarities between the reduction of long-range connections and short-range connections then we saw that young SC matrix much more similar to old SC matrix for reduction of long-range connections than that of short-range. But similarity gradually decreases as we deleted more and more connections. So, young SC became more like old SC as we deleted long-range connections, then we simulated the model to explore how metastability affected by long-range structural disruption.

**Figure 13:**
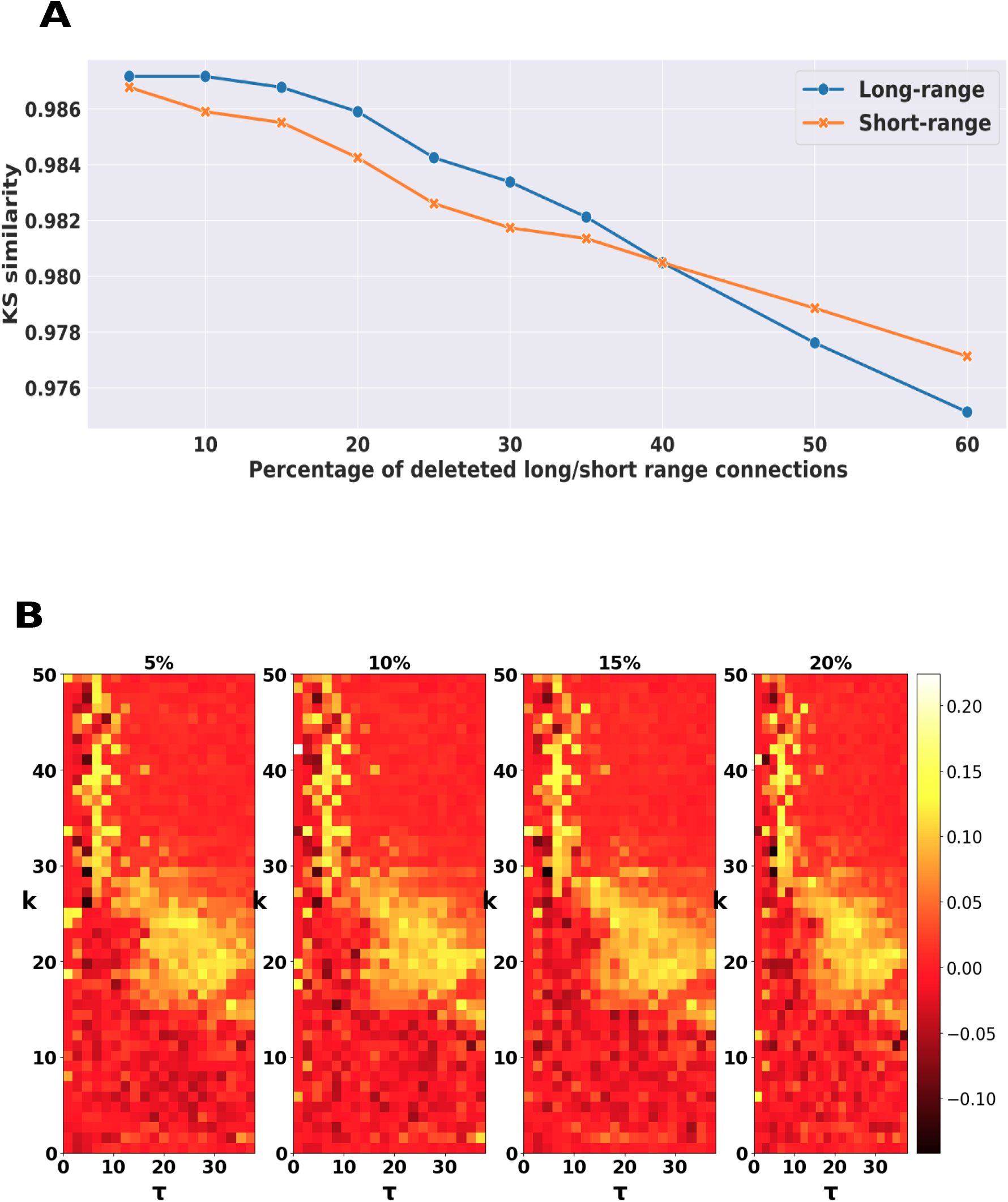
Similarity between long- and short-range connectivity profiles. (A) KS similarity of empirical structural connectivity of old group and synthetic structural connectivity matrix with various percentages of reduced long- and short-range connections. (B) Differences of metastability in (*τ, k*) parametric space for 5%, 10%, 20%, and 25% deleted long-range connections respectively. Difference is calculated by subtracting the metastability of young subject from the metastability of modified structural connectivity matrix.

To establish whether long-range connections has any direct relation with global metastability, we have deleted top 5% long-range structural connections from structural connectivity matrix of young subject and calculated the difference by subtracting the global metastability of young subjects from modified global metastability with deleted connections. So the positive difference will clearly indicate the enhancement of metastability after deletion of long-range connections. We repeat the steps for 10%, 15%, and 20% deleted connections. We plotted difference of global metastability in Fig. 13 (B) for 5%,10%, 15% and 20% deleted long-range connections respectively. Long-range connections that had been deleted from SC matrix were mainly long-range connections from the nodes of SN, CEN networks. Positive difference in parameter space indicates the global metastability increased as we deleted long-range connections.

## 4 Discussion

Neuroscience literature has primarily emphasized the investigation of metastability in models of complex behaviors, especially those that demand dynamic network interactions. So, we simulate a whole-brain Kuramoto model of coupled oscillators with appropriate conduction delay and interareal coupling strength to test the hypothesis of shifting of dynamic working point with age-associated alteration in network dynamics in both neural and ultraslow BOLD signal time scales by using synchrony and metastability [24, 49].

Aging entails variability involving different timescale and metastability may be consistent with other descriptions of capturing brain signal variability at slow and fast time scales. Naik et al [11] suggested to involve the concept of transient and metastability coordination to track fluidity of aging and cognition in fast and slow temporal scales. But it is unclear to what extent this metastability is able to capture this variability in different time scales. We have observed that the values of metastability at one timescale has no seemingly direct bearing on the metastability value at the other timescale. On the other hand, we can also observe that a higher (lower) neural activity across the BOLD timescale also corresponded to a similar higher (lower) neural activity at the neural timescale. It was thus exciting to note that despite the apparent similarity in the neural activity across the timescales, the metastability of the networks at the two timescales remains independent of each other. So, metastability can track the dynamics changes ranging from microscopic neural scale to macroscopic BOLD scales of the brain. Metastability measured using fMRI was enhanced in older subjects compared to younger subjects. Global and local metastability increased with age which is in line with previous such comparative studies [10, 11, 17]. Small and subtle changes in the brain across the lifetime of individuals might provide the basis for the increased metastability for the old population. In view of this, the increase in metastability for the old population present an interesting and rather counter intuitive case! It may be hypothesized that this increase may be an inherent coping mechanism of the brain that served to maintain an optimal degree of cognitive performance by keeping the brain more ‘ready’.

To understand the underlying inherent coping mechanism of the brain that served to maintain an optimal degree of cognitive performance, we simulated a simple biophysical model-the Kuramoto model-whose behavior has been rather well studied theoretically for generating neural activity [24, 31]. In this model, the circular phase is sufficient to represent the activity of a local system (neuron/neural column/cortical area). To make the model more neurobiologically plausible, time-delay version of the Kuramoto model with two ‘free’ parameters, namely the mean time delay, *τ* and the global coupling constant, *k* are introduced. Time delays in neuronal systems generated from finite axonal transmission, which is dependent on inter-areal distance and myelination as well as on synaptic and dendritic processes. Finally, the model was tuned by sweeping across this two-dimensional parameter space, keeping within bio-physically realistic limits, thus leading to the ‘parameter maps’ of measures such as synchronization, metastability, etc. which characterizes the relationship of the underlying anatomy with these network measures. We first calculated the simulated FC and compared it to the empirical one. We found a region where the simulated FC best fits the empirical one. The region of high correlation was confined to a much smaller area with smaller coupling strength in the case of the older population group. This reduction in the older population may be viewed as a case of the brain becoming more idiosyncratic with age, something which has been reported in literature [50]. In the young population, a region of high FC correlation suggested that the Kuramoto model can be a good source of generating realistic neural activity. At the same time, the reduction in correlation values for the old group indicated that as the brain ages, the model predicted parameter space may tend to shrink and shift offering lesser accuracy between model fit and the empirical observations.

The region where simulated FC best matches with empirical FC was delimited by an interval of sufficiently high coupling and realistic transmission delays where BOLD signals exhibited a significant level of partial synchrony and metastability. The metastability values were small outside this region for both group. This suggests in order to work brain at an optimal level some regions of brain were needed to be coupled as well as decoupled. Also, if metastability, as it is often described, is the measure of the preparedness of the brain to switch between tasks [51], the above results imply that this preparedness was highest when the brain was partially synchronized. With high metastability for old cohort, they were most likely to take less transitioning between distinct cognitive states at rest. In other words, metastability is, in this respect, a measure of the balance between the integration and segregation functionalities of the brain [22, 23], and thus the level of the metastability can be a marker for the optimum performance of the brain.

The optimal working region got shirked with smaller correlation value for older group of population but empirically global metastability increased for older population. So we can hypothesize that some local large scale networks are recruited for maintaining its high metastability. Metastability used for coexistence of compensation and dedifferentiation of different aging theories such as default to executive coupling hypothesis of aging also known as DECHA [52], posterior to anterior shift in aging (PASA). SN, involve in a variety of higher cognitive functions including communication, emotion processing etc. as well as being responsible for switching between the CEN and the DMN [41, 42] has higher metastability in the same optimal region for older population. Our computational findings, alongside empirical observations, provide additional support that Metastability helps to capture the fact that may be for healthy aging global brain acting optimally but there are some network scaffolds those are recruited for maintaining the optimal integrity of the brain with age.

To understand the increased relationship of metastability across topologically distinct functional networks over the lifespan, we explored the changes in underlying structural connectivity. The wiring pattern and structure of the brain have been the main focus of the field of connectomics and network neuroscience in general. The structure of the brain is not constant and changes to it occur during the lifespan of the individual. We know that the anatomical structure of the brain follows the so called ‘small-world’ organization [53], which has been found to be widespread in both artificial and natural systems [54, 55]. Structurally, age-related changes have shown a decrease in connection strengths [56] which may be a result of demyelination or loss of fibers. This loss in the structural consistency of the brain may be one of the main factors in the decline of cognitive ability in aging individuals, which is well documented even in the case of healthy aging. A key characteristic of brain networks is the distribution profile of the connection strength with the fiber distance. We saw that as the brain ages, there was a reduction in the number of connections between the different regions of the brain. However, the slopes of the SC strength versus the fiber length distribution showed no significant change across the two age groups. The maintaining of this distribution profile perhaps may be one of the keys to healthy aging as opposed to pathological aging.

We were more interested to check whether the long-range and short-range connections could be the reason for higher metastability. In general long-range and short-range tracts are another important elements of brain structure that are of key importance for the functional-anatomical organization of the cortex. The higher cognitive functional networks are not preferentially limited to isolated cortical areas rather widespread over the brain. This leads to the conclusion that the long-range connections between cortical and sub-cortical areas must be a fundamental structural parameter in order to sustain the functional complexity [57]. So the long-range connections, bring down the inter-areal distances, are a strong factor for efficient communication between functional networks [58].

We found that number of long-range white-matter fiber connections reduced significantly with age which is in line with the previous findings [7].The disruptions in long-range connections lower the involvement of number of brain regions in the different functional modules [59] yielding the fact that the aging brain needs more steps via short-range connections to transfer information from one part of the brain to another. So we can hypothesize that the brain creates more small clusters by coupling more regions to maintain its optimal performance. This encourages partial synchronization with metastability to enhance to compensate for the structural damages. Naik et al. [11] also suggested the same that high-functioning adults would exhibit higher metastability to compensate for the structural damages of brain networks during senescence. So, a lesser number of long-range connections could give rise to higher metastability in older individuals. We analytically had shown that indirect anatomical links have contribution to synchronization manifold. We numerically has proved that reduced long-range connectivity in local resting state networks give rise to metastability in global level.

Although we could convincingly demonstrate analytically and numerically how structural connectomics plays a crucial role in determining age-associated alterations in network synchrony and metastable brain dynamics both at neuronal and BOLD time scales, there are also several limitations to this work. The Kuramoto model, a system of coupled oscillators, is a less complex generalization of brain function. For example, the simulation was built on a relatively low-dimensional connectivity matrix of 68 regions. The sample size was also moderate, and we may have restricted power to detect some effect, particularly in structural connection to network dynamics. In addition, there are certain limitations inherent in tractography measured with diffusion MR. It may be challenging to resolve accurately as uncertainty in streamlined location increases with the length of the tract. However, despite these limitations, our whole brain model shed crucial insight into the nature of the relationship between structural connectomics with age and invariance of network measures synchrony and metastability operating on both time scales of cognition. This is broadly consistent with empirical findings. We plan to extend the network-level analysis with a larger group of subjects to correlate the network-level findings with different cognitive processes. Another future direction could be to consider whether these results sufficiently generalize while employing other neural mass models, such as the Wilson-Cowan model. Furthermore, various graph theoretical measures, such as clustering coefficient and short path length and network communication models such as spreading cascade, could provide additional insights for Spatio-temporal alterations in metastability and transient waves that could be examined in future brain network studies.

